# Development of a high-throughput homogeneous AlphaLISA drug screening assay for the detection of SARS-CoV-2 Nucleocapsid

**DOI:** 10.1101/2020.08.20.258129

**Authors:** Kirill Gorshkov, Catherine Z. Chen, Juan Carlos de la Torre, Luis Martinez-Sobrido, Thomas Moran, Wei Zheng

## Abstract

The coronavirus disease 2019 (COVID-19) pandemic caused by Severe Acute Respiratory Syndrome coronavirus 2 (SARS-CoV-2) is in urgent need of therapeutic options. High-throughput screening (HTS) offers the research field an opportunity to rapidly identify such compounds. In this work, we have developed a homogeneous cell-based HTS system using AlphaLISA detection technology for the SARS-CoV-2 nucleocapsid protein (NP). Our assay measures both recombinant NP and endogenous NP from viral lysates and tissue culture supernatants (TCS) in a sandwich-based format using two monoclonal antibodies against the NP analyte. Viral NP was detected and quantified in both tissue culture supernatants and cell lysates, with large differences observed between 24 hours and 48 hours of infection. We simulated the viral infection by spiking in recombinant NP into 384-well plates with live Vero-E6 cells and were able to detect the NP with high sensitivity and a large dynamic range. Anti-viral agents that inhibit either viral cell entry or replication will decrease the AlphaLISA NP signal. Thus, this assay can be used for high-throughput screening of small molecules and biologics in the fight against the COVID-19 pandemic.

## Introduction

The <u>co</u>rona<u>vi</u>rus <u>d</u>isease of 20<u>19</u>, COVID-19, has spread rapidly through the global population since late 2019 due to the newly highly infectious and deadly <u>s</u>udden <u>a</u>cute <u>r</u>espiratory <u>s</u>yndrome <u>co</u>rona<u>v</u>irus 2, SARS-CoV-2^1^. The unprecedented response from the global research community at every level of the life science enterprise has produced several therapeutic candidates such as the repurposed investigational Ebola drug remdesivir that targets the SARS-CoV-2 RNA-dependent RNA polymerase ^2,3^, neutralizing antibodies against the viral spike (S) protein^4^, and convalescent plasma from recovered patients containing neutralizing antibodies^5^. Additionally, hundreds of vaccines against SARS-CoV-2 are currently under development with over two dozens of them in clinical phase I and II trials^6,7^. While these are all viable options, no therapeutic option is yet approved by the United States Food and Drug Administration (FDA). Therefore, we must continue the search for other alternatives, a task that will benefit from new assays and reagents that facilitate rapid identification of small molecules and biologics that suppress viral infection to treat COVID-19.

High-throughput screening (HTS) is one such method wherein high density microplates can be used to rapidly screen large compound libraries to discover lead compounds for drug development^8^. It also can be used for drug repurposing screens and evaluating therapeutic agents with high confidence, low variability, and generate information regarding an agent’s potency via half-maximal inhibitory concentration (IC_50_) and its maximum efficacy. Over the last decade, amplified luminescent proximity homogeneous assay (AlphaLISA) technology has emerged as one of the most reliable screening technologies because of its versatility, sensitivity, and homogeneous format without the need for plate evacuation or washing^9^. AlphaLISA signal is generated when a donor bead is excited at an appropriate wavelength to general a reactive singlet oxygen radical that, when in close proximity to an acceptor bead, activates the acceptor bead to produce light at a specific wavelength^9^. It can be used to replace ELISA that requires multiple plate-wash steps for the quantitative detection of analytes. The conjugation of antibodies to biotin and streptavidin aids in the creation of a screening system that can be used as a competitive assay or sandwich assay format. In a competitive assay that uses one antibody, unlabeled analyte would compete with a labeled analyte donor or acceptor and decrease signal. In a sandwich assay, two antibodies raised against the same immunogen analyte, albeit at different epitopes, will both bind to the analyte and lead to an increase in signal.

The SARS-CoV-2 virion is made of four main component structural proteins including S, Envelope (E), Nucleocapsid (NP), and Membrane (M)^10^. SARS-CoV-2 S is responsible for initial host-cell binding to the angiotensin converting enzyme 2 (ACE2) receptor via the receptor binding domain (RBD) found within the S1 subunit^11^. E protein is embedded in the viral membrane and plays a role in viral particle packaging and maturation^12^. M protein regulates replication and RNA packaging^10^. The NP binds M-protein and interacts with the viral genome to bind the positive strand RNA viral genome, and it is the main focus of the work presented here^10^.

Here, we describe the development of a high-throughput SARS-CoV-2 NP-based immunoassay in the AlphaLISA sandwich assay format that can be used to detect live virus infection and replication in host cells to carry out HTS for drug discovery and development. We show the selection process for identifying the most reliable antibody pair to detect NP, along with the optimization of the AlphaLISA reagents for the highest sensitivity and dynamic range using recombinant NP. We further demonstrate this assay can detect the SARS-CoV-2 NP in cell lysates and tissue culture supernatants (TCS) after SARS-CoV-2 infection. Lastly, we have simulated the viral infection assay by spiking in recombinant NP into Vero-E6 cells grown in in 384-well plates, demonstrating the assay potential for HTS. The results presented here demonstrated a rapid, homogeneous SARS-CoV-2 NP assay that can be used for HTS of compound collections to identify compounds that inhibit SARS-CoV-2 infection and viral replication in live cells.

## Results

### Best NP antibody pair selection

The NP is one of the four structural proteins present in the SARS-CoV-2 particle, and its levels increase with viral replication as more viral particles are produced. Thus, the level of NP can serve as an indicator of virus infection and replication in host cells. The cell-based AlphaLISA SARS-CoV-2 NP detection assay relies on the proximity of two labeled antibodies that bind to NPs in cell lysate and supernatant in high-density microplates to generate the luminescent signal. This assay detects viral proteins inside the cell, and the virus that is released into the medium.

The biotinylated antibody is bound to the streptavidin coated donor bead that when excited with 680 nm light, induces a singlet oxygen radical to activate an acceptor bead conjugated to the second antibody, resulting in a 615 nm luminescent signal (Fig. 1A). A successful reaction depends on the close proximity of the donor and acceptor. Multiple antibodies were matrixed together to determine which antibody pair produced the highest AlphaLISA signal (counts) from recombinant NP at 10,000 pg/mL (Fig. 1B,C). The three best antibody pairs were 1+4, 11+4, and 11+10. Each pair was then tested against a titration of recombinant NP at 10,000 pg/mL, 1000 pg/mL, 100 pg/mL and 0 pg/mL, and produced a concentration dependent signal increase, with signal to background ratios (S/B, 10,000 pg/mL to 0 mg/mL) of 66.2, 72.8, and 51.5 based on the S/B, respectively (Supplementary Fig. 1). Based on these results, pairs 1+4, and 11+4 were selected for further optimization. Interestingly, the pairs consisting of antibody 11 did not detect the Histidine (His)-tagged NP (Fig. 1B), but generated the best signal with untagged NP (Fig. 1C), suggesting the His-tag interferes with binding of antibody 11 to NP. The other antibody pairs produced relatively similar counts regardless of the NP protein used in the optimization.

**Figure 1.**
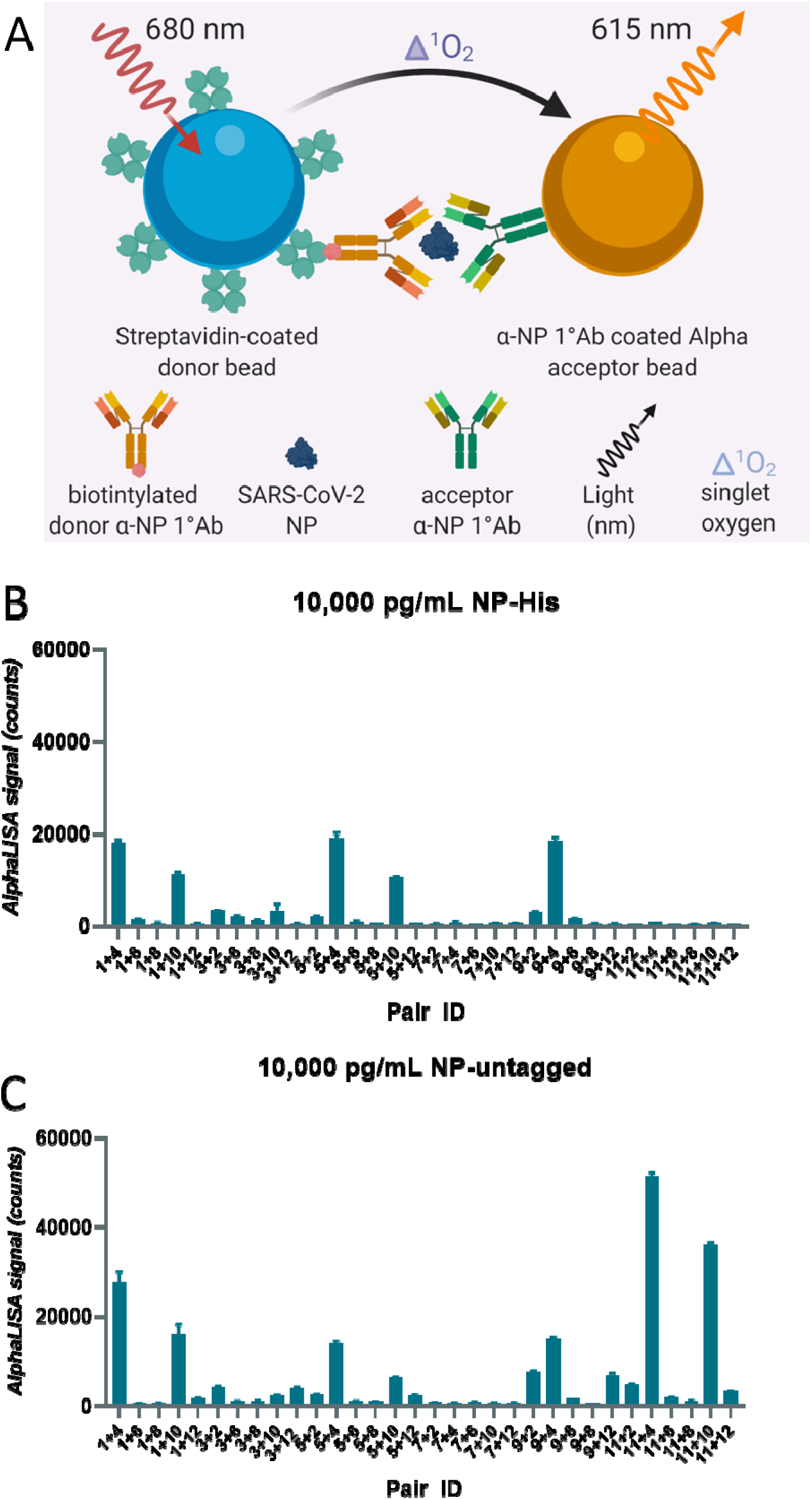
Best pair antibody selection for SARS-CoV-2 Nucleocapsid AlphaLISA sandwich assay. **(A)** Schematic diagram of homogenous cell-based AlphaLISA sandwich assay. **(B)** Table of matrixed antibodies and the concentrations of recombinant SARS-CoV-2 Nucleocapsid. **(C)** AlphaLISA signal (counts) for matrixed antibody pairs using Nucleocapsid-His tagged. High values indicate efficient AlphaLISA reaction. **(D)** AlphaLISA signal (counts) for matrixed antibody pairs using untagged Nucleocapsid. High values indicate efficient AlphaLISA reaction. N = duplicate wells. Error bars indicate S.D.

### Optimization of acceptor to donor ratio and biotin concentration

We next determined the S/B at a greater range of untagged recombinant NP (20,000 pg/mL – 81.9 pg/mL) using a donor/SA acceptor ratio of 10 μg/mL to 40 μg/mL and 20 μg/mL to 20 μg/mL and different concentrations of biotin from 0.5 nM to 5.0 nM (Supplementary Fig. 2A-D). We selected a donor/SA ratio of 10 μg/mL to 40 μg/mL for both pairs, and 1.0 nM and 5.0 nM of biotin for Pairs 1+4 and 11+4, respectively, based on the S/B calculations (Supplementary Fig. 2D). The standard curves were generated using recombinant NP for these conditions and no plateau was observed at the higher concentrations, indicating that the upper limit of detection was more than 20,000 pg/mL (20 ng/mL).

### Lysis buffer selection and viral lysate testing

To determine the limits of detection and dynamic range of the assay, we next generated standard curves using concentrations of NP from 200,000 ng/mL to 0.02 ng/mL for both pairs (Fig. 2A,E). We observed a significant hook effect at concentrations of recombinant NP above 320 ng/mL for pair 1+4, and 1600 ng/mL for pair 11+4 (Fig. 2B,F). The hook effect indicates the concentration of analyte at which an overabundance of analyte titrates the donor and acceptor away from each other, and this hooking dictates the upper limit of detection before the signal will start to decrease. Two different lysis buffers were used to optimize any lysis buffer effects; the NCATS lysis buffer consisted of 0.5% Triton-X 100 with protease inhibitor, the other was the AlphaLISA lysis buffer from the manufacturer. For pair 1+4, both lysis buffers performed equally, while the AlphaLISA lysis buffer performed better for pair 11+4 in generating the standard curve. The overall S/B was higher for pair 11+4 at higher concentrations of NP, but pair 1+4 had a higher sensitivity as seen by the larger S/B values at lower concentrations (Fig. 2B,F).

**Figure 2.**
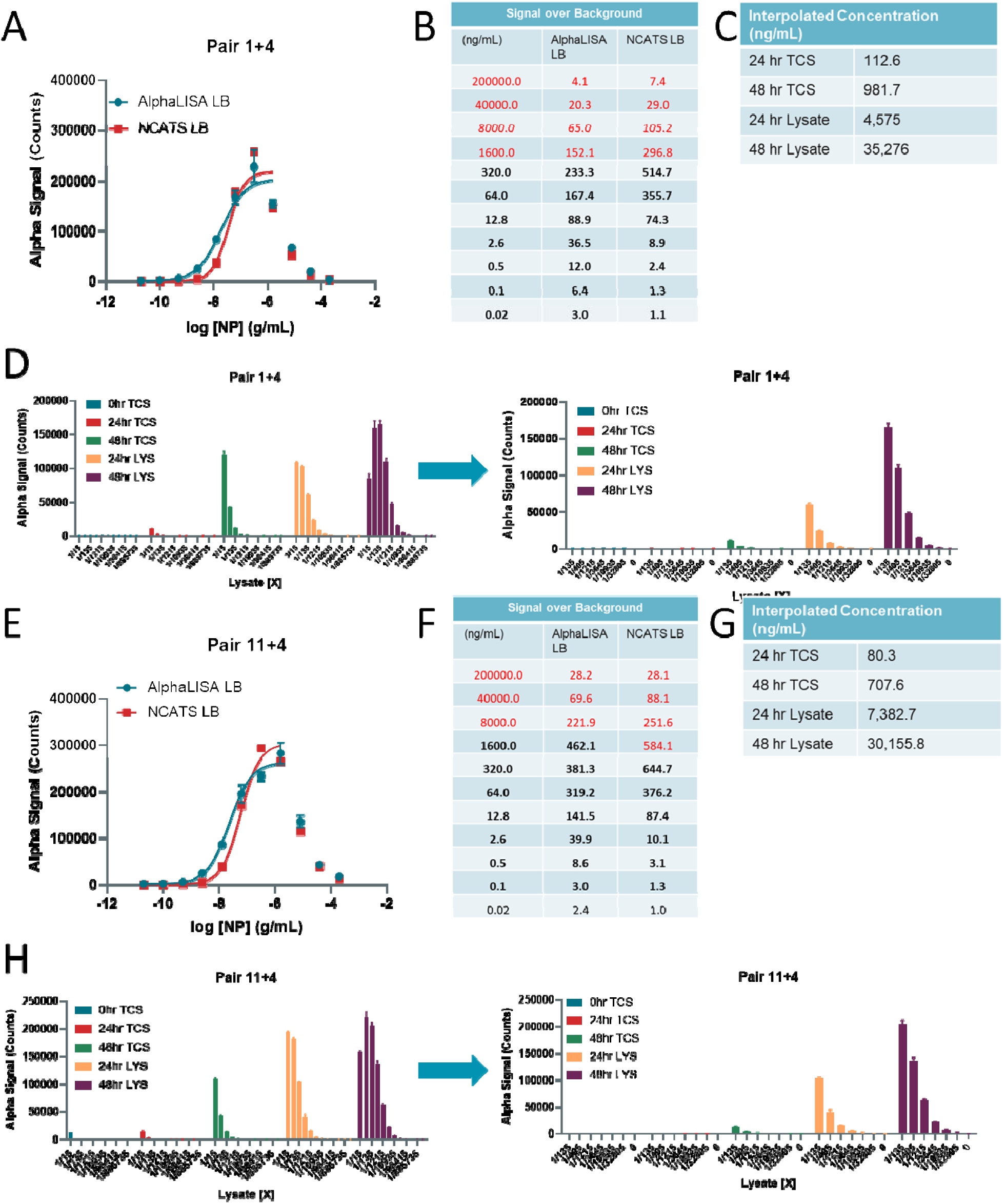
SARS-CoV-2 Nucleocapsid detection in virally infected cell lysates and tissue culture supernatants. **(A)** Standard curve for Pair 1+4 demonstrating hook effect using AlphaLISA or NCATS lysis buffer. **(B)** Signal over Background table for A. Values in red show the concentrations at which accurate determination of Nucloecapsid is not possible due to the hook effect. **(C)** (Left) AlphaLISA signal (counts) for 24 hour and 48 hour virally infected cell lysates and 0 hour, 24 hour, and 48 hour tissue culture supernatants at various dilutions. (Right) Same as left panel with a narrower dilution range. **(D)** Standard curve for Pair 11+4 demonstrating hook effect using AlphaLisA or NCATS lysis buffer. **(E)** Signal over Background table for D. Values in red show the concentrations at which accurate determination of Nucloecapsid is not possible due to the hook effect. **(F)** (Left) AlphaLISA signal (counts) for 24 hour and 48 hour virally infected cell lysates and 0 hour, 24 hour, and 48 hour tissue culture supernatants at various dilutions. (Right) Same as left panel with a narrower dilution range. N = triplicate wells. Error bars indicate S.D.

To determine whether the detection system could detect native NP in SARS-CoV-2 infected lysates, TCS collected at the 0 hr, 24 hr, and 48 hr TCS along with cell lysates collected at 24 and 48 hr post-infection were tested at multiple dilutions. Pair 11+4 produced greater absolute AlphaLISA signal (counts) than pair 1+4 at the highest concentrations, but both pairs exhibited a hook effect at 1/15 lysate dilutions (Fig. 2D,H). NP was detected in TCS by pair 11+4 better than pair 1+4. The concentrations of NP in ng/mL were interpolated from the standard curves. For both pairs, the concentration of NP was approximately 9-fold greater in 48 hr TCS than 24 hr TCS. For pair 11+4, the 24 hr 48 hr TCS was 80.3 ng/mL and 708 ng/mL, respectively. The 24 hr and 48 hr lysate was significantly higher at 7,380 ng/mL and 30,200 ng/mL, respectively. These concentrations were comparable, although not exactly the same, when comparing pair 1+4 and 11+4.

### Sequential two-step v.s. three-step assay protocol optimization

We next optimized the sequential addition of assay reagents including lysis buffer, biotin, acceptor beads, and donor beads in the two-step v.s. three-step assay protocol (Supplementary Fig. 4A,B). For pairs 1+4 and 11+4, the two-step assay produced a greater dynamic range for recombinant NP detection than the three-step assay without exhibiting a hook effect (Supplementary Fig. 4C-J) as determined by the S/B ratios. NCATS lysis buffer performed slightly better than AlphaLISA lysis buffer. We further optimized the two-step and three-step assays for recombinant NP spiked into Vero-E6 cells grown in 384-well plates (Supplementary Fig. 5). In most cases, the two-step assays performed better with 20,000 cells per well as measured by S/B ratios. The effect of media and cell density was also tested for assay interference. Media affected pair 1+4 to a greater extent than pair 11+4, while cell density did not significantly affect S/B ratios. The lower limit of detection for both pairs was 0.22 ng/mL with an upper limit at or beyond 400 ng/mL (Supplementary Fig. 6).

### Assay volume optimization

Next, we optimized the total assay volume in 384-well plate format using recombinant NP in AlphaLISA LB and found that higher total assay volumes produced lower absolute AlphaLISA signal (counts), but 100 uL total volume performed better than 50 uL total volume (Fig. 3A,C). Again, pair 11+4 exhibited a higher limit of detection than pair 1+4 (Fig. 3B,D). In this assay, the 20 μL condition utilized the 384-well Proxiplate, while the 50 uL and 100 uL conditions utilized the 384-well CulturPlate. The main difference between the two is the shallower circle wells in the Proxiplate compared to the deeper square wells in the CulturPlate, suggesting the plate type and well dimensions does change the S/B ratios.

**Figure 3.**
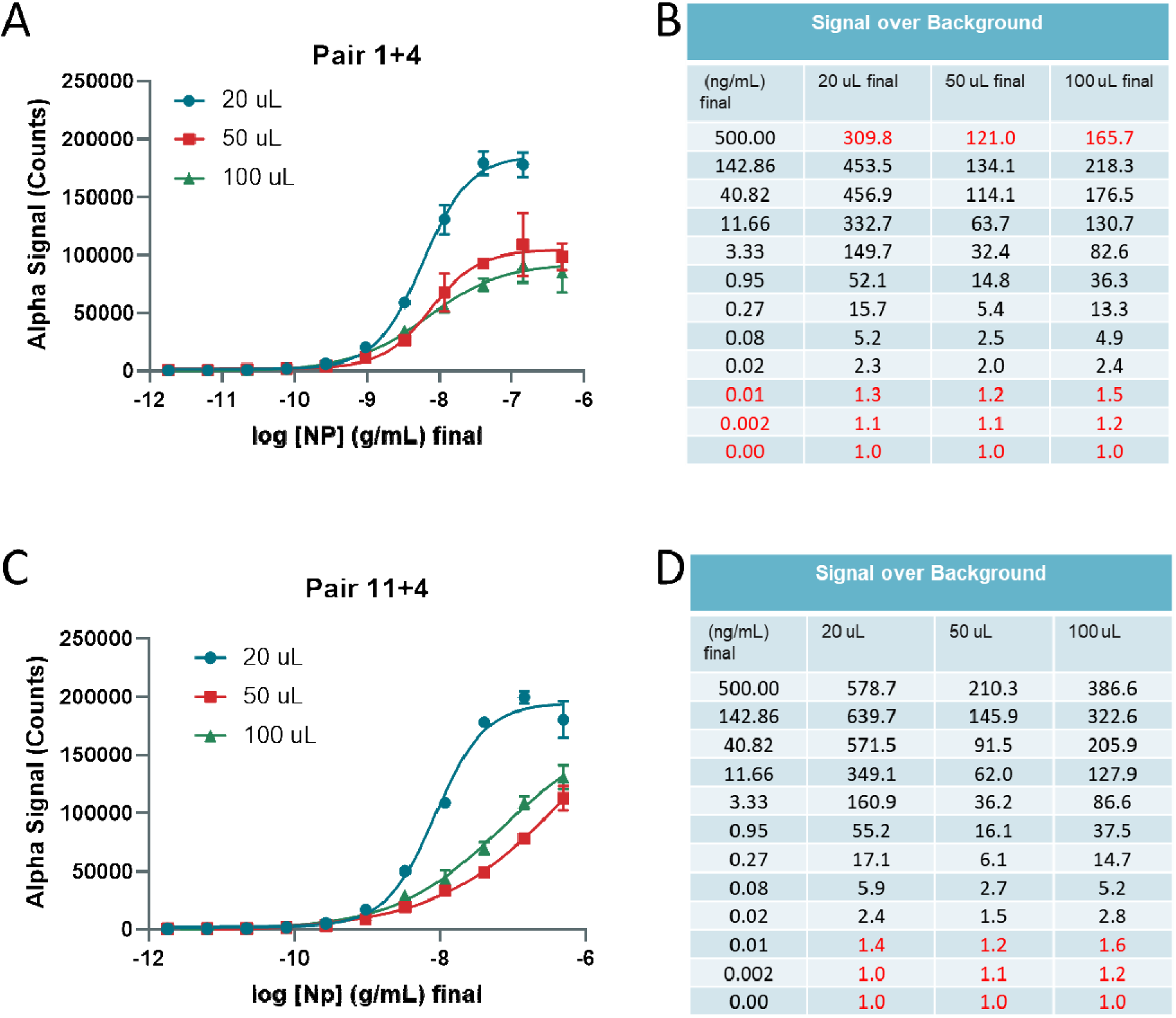
Optimization of assay volumes for 384-well plate using recombinant Nucleocapsid. **(A)** Detection of titrated Nucleocapsid in wells with different final volumes of 20, 50 and 100 μL using Pair 1+4. **(B)** Signal over Background table from A. Red values indicate hook effect (upper limit) and lower limit of detection. **(A)** Detection of titrated Nucleocapsid in wells with different starting volumes 20, 50 and 100 μL using Pair 11+4. **(B)** Signal over Background table from A. Red values indicate lower limit of detection. N = triplicate wells. Error bars indicate S.D.

We further optimized media conditions and found that cell culture media itself had a profound effect on the S/B ratios, while the concentration of FBS had little to no impact on the assay performance (Supplementary Fig. 7).

### Batch to batch reagent reproducibility

A second batch of reagents was produced and the concentration of biotin was matched to generate the same performance as the first batch (for pair 1+4 and pair 11+4 Supplementary Fig. 8). We tested the second batch of reagents with the recombinant NP spiked into Vero-E6 cell culture in 384-well plates and found a better performance with pair 11+4 when cells were present in the wells (Fig. 4A-D). The lower limit of detection of 0.2 ng/mL and an upper limit of greater than 500 ng/mL NP. Altogether, the pair 11+4 performed better than pair 1+4 and was selected as the final pair for future assays. To confirm that the second batch of reagents performed well, we tested the same conditions for optimization as before with cell densities ranging from 20,000 cells per well to 0 cells per well, along with a no media control in PBS.

**Figure 4.**
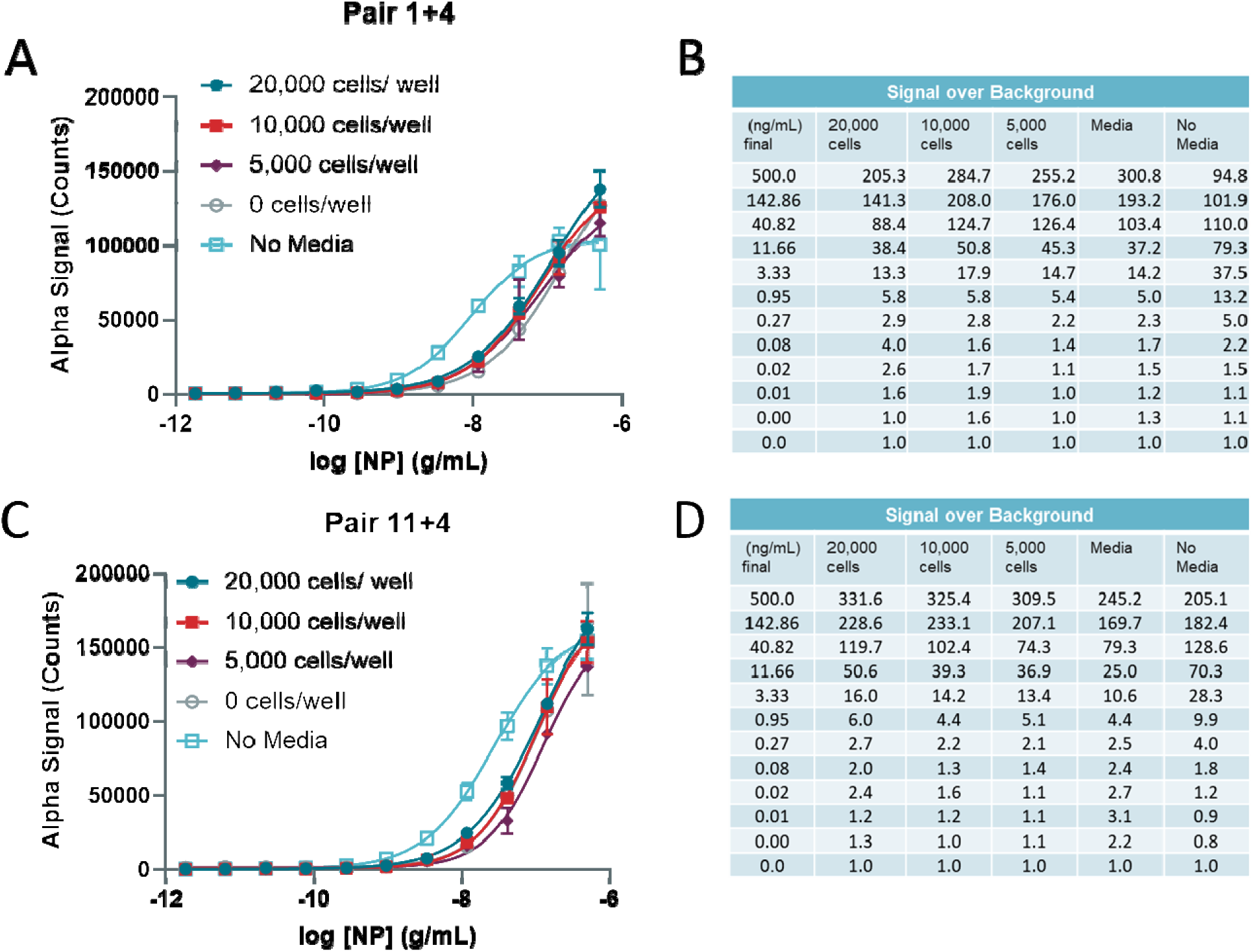
Optimization of 384-well plate cell number in simulated infection using second preparation of reagents. **(A)** Detection of titrated Nucleocapsid in wells culturing 20,000, 10,000, 5000, or 0 cells/well along with a no-media control in Vero-E6 cells using Pair 1+4 (second batch). **(B)** Signal over Background table from C. **(C)** Detection of titrated Nucleocapsid in wells culturing 20,000, 10,000, 5000, or 0 cells/well along with a no-media control in Vero-E6 cells using Pair 11+4 (second batch). **(D)** Signal over Background table from E. N = triplicate wells. Error bars indicate S.D.

### Lysis buffer and incubation time optimization, and determination of cell lysate interference

The AlphaLISA buffer and NCATS lysis buffers (LB) were compared in plates incubated for 30 min. v.s. 1 hr and for the AlphaLISA buffer there was a slight increase in signal after 1 hour incubation (Supplementary Fig. 9A-D). The incubation time had little effect on the S/B for the NCATS LB condition. NP detection in the presence of lysed 10,000 and 5,000 cell per well performed similarly to NP detection in media, and better than in 20,000 cells per well. 20,000 cells per well would be considered over-confluent beyond 100%. The no media PBS control had a lower S/B ratio overall. The main difference between the LBs was that a signal plateau was observed for the NCATS LB indicating an upper limit of detection of approximately 1500 ng/mL, whereas the AlphaLISA buffer could potentially detect higher concentrations. Importantly, the sensitivity of the assay with NCATS LB reveals a similar lower limit of detection of 0.23 ng/mL as in previous experiments, whereas the AlphaLISA buffer was observed be less sensitive with a lower limit of detection of 2.06 ng/mL for 10,000 cells per well, for example.

### Estimation of NP concentration in cell lysates using standard curve interpolation

Finally, we estimated the concentration of NP in TCS and lysate from cells infected with live SARS-CoV-2 for 24 hr and 48 hr compared to the non-infected (0 hr) controls using interpolation from the 10,000 cells per well standard curve in Supplementary Figure 9E (Supplementary Fig. 10A-C). Dilutions of TCS and lysates from 15-fold to over 1900-fold were created and NCATS LB was added to the samples. The linear portion of the curves was used for interpolation. The concentrations of NP in lysates was determined to be 7,300 and 34,092 ng/mL for 24 hr and 48 hr lysates, respectively (Supplementary Fig. 10C). In correspondence with the NP concentration, the viral titers were calculated to be 8.2E+06 focus forming units (FFU)/mL and 1.1E+09 FFU/mL for 24 hr and 48 hr TCS, respectively, indicating increasing amounts of viral particles with longer infection times. This data indicated that the NP in viral lysates could be quantified using the AlphaLISA NP detection reagents and standard curve interpolation.

## Discussion

The ability to rapidly detect viral proteins in a homogeneous cell culture system provides researchers with an easy to use method for screening thousands of potential antiviral compounds, both newly synthesized analogs and existing FDA-approved drugs alike. In this work we have designed and developed through meticulous optimization an AlphaLISA sandwich-based detection system to identify the SARS-CoV-2 NP. Here, two antibodies were identified to detect recombinant and virally-generated NP both in solution and in cell culture. Importantly, we demonstrated a significant increase in NP with longer incubation of virally-infected Vero-E6 cells and were able to quantitatively assess the NP concentration using standard curve interpolation. This assay system should be able to identify compounds that decrease the presence of SARS-CoV-2 NP as a measure of anti-viral activity that either prevents viral entry or viral replication leading to the translation of NP in cells. Our add-and-read system requires no wash steps or removal of cell culture media, providing an excellent methodology for the use in biosafety level 3/4 (BSL 3/4) facilities that require simple and easy to use assay technologies for drug screening and validation.

The selection of SARS-CoV-2 NP was a strategic one in that mutations of the SARS-CoV-2 viral genome will likely occur in coding regions that are selected for during viral-host interactions. The SARS-CoV-2 viral S protein that protrudes from the envelope binds ACE2 with high affinity as the first step in viral infection^13^, whereas the viral NP is located deeper within the virion core. Indeed, mutations in Spike have already been identified, with one of those being the more infectious D614G mutation^14–16^. By targeting NP, this detection system will maintain its reliability for drug screening of multiple viral strains, barring significant changes to the epitopes to which the antibodies used in the AlphaLISA sandwich assay bind.

We anticipate rapid developments in HTS screening following the adoption and application of the AlphaLISA NP detection system in BSL 3/4 laboratories. Future work will optimize assay conditions for live virus experiments to establish the appropriate multiplicity of infection and incubation time of the virus in order to match the wide dynamic range and high sensitivity of the assay.

## Methods

### Reagents and Materials

The following item was purchased from ATCC: Vero-E6 (CRL-1586, RRID:CVCL_0574). The following items were purchased from Corning TM: EMEM (10-009-CV), HI FBS (35-016-VC) and 0.25% Trypsin (25053CI). The untagged NP (Z03501) was purchased from Genscript. The His-tagged NP (40588-V08B) was purchased from SinoBiological.

The following item was purchased from Gibco: Pen/Strep (15140-122). PBS (SH30256FS) was purchased from HyClone. The following items were purchased from PerkinElmer: ProxiPlate-384 Plus (Cat# 6008280), CulturPlate-384 (Cat#: 6007680), Alpha Streptavidin Donor beads (6760002), AlphaLISA lysis buffer (AL003C). The following custom labeling was performed by PerkinElmer: donor antibodies were biotinylated using NHS activated biotinylating reagent (ChromaLink #B-1001-105), and acceptor antibodies were conjugated to Alpha Acceptor Beads (PerkinElmer #6760137M).

### Preparation of antibody pair matrixing

A matrix with all antibodies provided by PerkinElmer was tested for the ability to detect NP.

The untagged NP and His-tagged NP were diluted in AlphaLISA lysis buffer at 10,000; 1,000 and 100 pg/mL. For initial testing, 20 μg/mL AlphaLISA Acceptor (final), 1 nM (final) biotinylated antibody and 20 μg/mL Donor (final) were used. A two-step assay was performed (data points in duplicate) using the following protocol:

1. Dispense 5 μl of recombinant protein.
2. Dispense 10 μl of mix of acceptor beads and biotinylated antibody.
3. Incubate 60 min. at room temperature (RT).
4. Dispense 5 μL of SA-Donor Beads.
5. Incubate 30 min. at RT.
6. Read plates on EnVision (Perkin Elmer).

### Antibody concentration optimizations

Best pair 1+4 and 11+4 were optimized using the following parameters: Biotinylated antibody concentration were tested at 0.5 nM, 1 nM, 2 nM, and 5 nM. Acceptor bead – SA donor beads concentrations were tested at 20 μg/mL – 20 μg/mL and 10 μg/mL – 40 μg/mL. Concentration of untagged NP recombinant protein started at 20,000 pg/mL followed by 2.5-fold dilutions. The dispensing protocol was the same as for the antibody pair matrixing.

### Vero-E6 cell culture

Vero-E6 (grown in EMEM, 10% FBS, and 1% Penicillin/Streptomycin), were cultured in T175 flasks and passaged at 95% confluency. Briefly, cells were washed once with PBS and dissociated from the flask using 0.25% Trypsin. Cells were counted prior to seeding.

### Preparation of viral lysate and tissue culture supernatant production

Vero-E6 cells were plated in 12-well plates at 450k cells/well in 1.25 mL of growth media. Cells were incubated for 24 hr at 37°C. 250 μL of SARS-CoV-2 (Washington Strain) was used at an MOI of 0.05. Cells were inoculated for 45 min. at 37°C. Supernatant was collected and pooled for 0 hr TCS. A final concentration of 0.5% Triton-X 100 and 1x protease inhibitor was used for all samples. TCS samples were stored at −20°C until needed. 1.5 mL of fresh media was added to wells and cells were incubated for 24 hr and 48 hr at 37°C. 6 wells were harvested at each time point. Supernatant was collected from 6 wells each at 24 hr and 48 hr. Samples were pooled for 24 hr or 48 hr TCS. 200 μL of TCS was removed for viral titer calculations and diluted with equal volume of lysis buffer. Samples were stored at −20°C until needed. Lysate was collected from 6 wells each at 24 hr and 48 hr. Cells were rinsed once with ice cold PBS. 250 μL/well of cell lysis buffer + PI was added to lyse cells. Protease inhibitor was added to the lysis buffer before lysing cells. Cells were scraped to the bottom of the well and pooled lysates were collected into 1.5 mL tubes on ice. Tubes were vortexed for 10 s and placed on ice for 10 min. This was repeated three times. Lysates were spun down for 30 min at 13.2k rpm at 4°C and stored at −20°C until further use.

### Optimization of assay using viral lysates and tissue culture supernatant

The best conditions of pairs 1+4 and 11+4 were used to expand the standard curve and test TCS and lysate samples:1+4: 10-40 μg/mL with 1 nM Biotin, 11+4: 10-40 μg/mL with 5 nM Biotin. An 11-point curve of recombinant untagged NP started at 200,000 ng/mL followed by 5-fold dilutions. Lysates and TCS were diluted in PBS + Triton-X 100 + protease inhibitors 1:15 (1X) followed by 3-fold dilutions. The dispensing protocol was the same as above.

### Simulation of viral infection using recombinant NP

The best conditions of pairs 1+4 and 11+4 were used to test the assay in a cell-based format: 1+4: 10-40 μg/mL with 1 nM Biotinylated antibody and 11+4: 10-40 μg/mL with 5 nM Biotinylated antibody. Vero E6 cells were plated at 20,000 and 50,000 cells/well in 384-well plate (TC) and incubated overnight. An 11-point curve of recombinant NP started at 4,000 ng/mL initial (400 ng/mL final) followed by 3.5-fold dilutions in AlphaLISA lysis buffer.

For second batch preparation, the best conditions of pairs 1+4 and 11+4 were used to test the assay in a cell-based format (newly conjugated and biotinylated Ab were used and optimized to reproduce previous antibodies): 1+4: 10-40 μg/ml with 0.5 nM Biotinylated antibody, 11+4: 10-40 μg/ml with 1 nM Biotinylated antibody. Vero E6 cells were plated at 20,000, 10,000, and 5,000 cells/well in 384w plate (TC) and incubated overnight. The assay was also tested without cells and without media as a control. An 11-point curve of recombinant NP started at 2,000 ng/mL initial (500 ng/mL final) followed by 3.5-fold dilutions in media.

### Statistical analysis and illustrations

Concentration-response curves were fit using non-linear regression, standard curve interpolation, and graphs were generated in Graphpad Prism V8.43. Illustration in Figure 1A was created using Biorender.

## Acknowledgements

This research was supported in part by the Intramural Research Program of the National Center for Advancing Translational Sciences, NIH. We thank Dr. Arturo Gonzalez-Moya and his team at Perkin Elmer for assay development support.

## Data Availability Statement

Data available upon request.

## Author contributions

Experimental contributions: KG, CC, JCT, LM-S,

TM Project management: KG, CC

Initial conception and design: KG, CC, WZ

Manuscript writing and editing: KG, CC, WZ

