## Supplemental figures for "Development of a high-throughput homogeneous AlphaLISA drug screening assay for the detection of SARS-CoV-2 Nucleocapsid"

**Supplementary Materials**

**Supplementary Figures and Figure Legends**

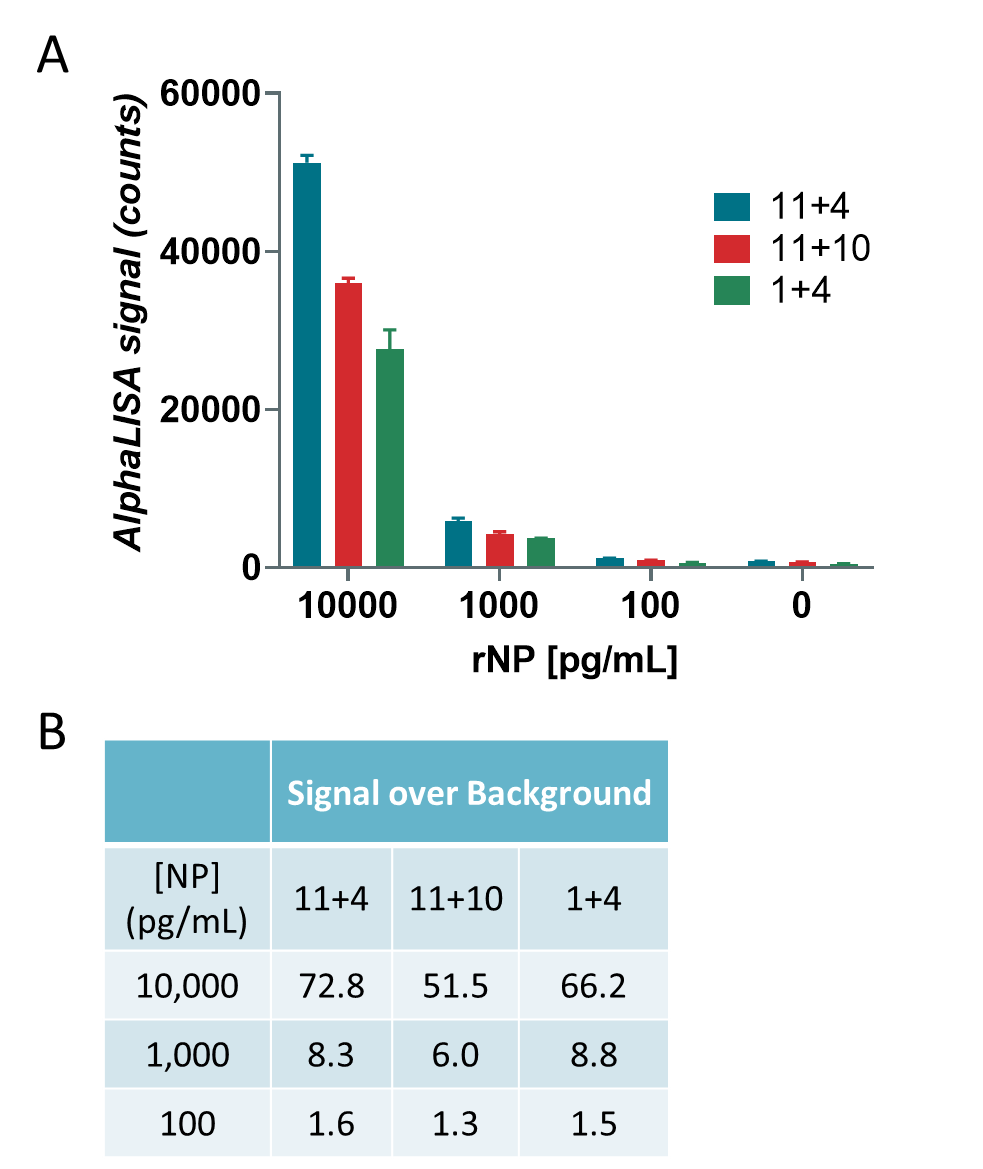

**Supplementary Figure 1.** Detection of titrated NP using best antibody pairs. AlphaLISA counts for **(A)** Pair 1+4, **(B)** Pair 11+4, **(C)** Pair 11+10 using 10,000, 1000, 100, and 0 pg/mL of NP. N = duplicate wells. Error bars indicate S.D.

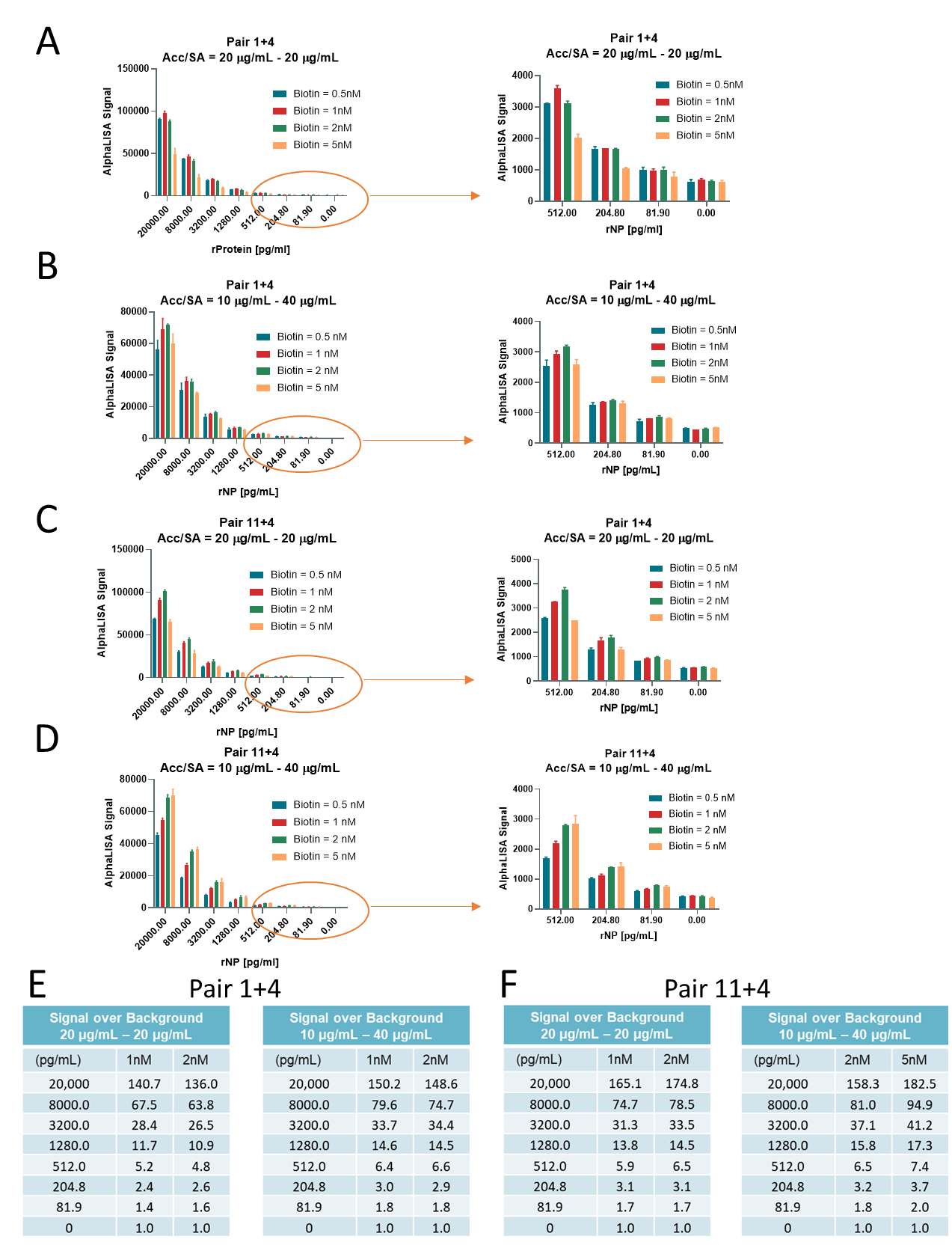

**Supplementary Figure 2.** Optimization of best antibody pair and reagent concentrations. AlphaLISA counts for titrated NP using **(A)** 20 µg/mL acceptor and 20 µg/mL or **(B)** 10 µg/mL acceptor and 40 µg/mL Streptavidin for Pair 1+4 and different concentrations of Biotin. Right panel is scaled section of left panel inside of red circle. AlphaLISA counts for titrated NP using **(C)** 20 µg/mL acceptor and 20 µg/mL or **(D)** 10 µg/mL acceptor and 40 µg/mL Streptavidin for Pair 11+4 and different concentrations of Biotin. Right panel is scaled section of left panel inside of red circle. **(E)** Table for graphs A and B using 1 nM and 2 nM Biotin. **(F)** Table for graphs C and D using 1 nM and 2 nM Biotin. N = duplicate wells. Error bars indicate S.D.

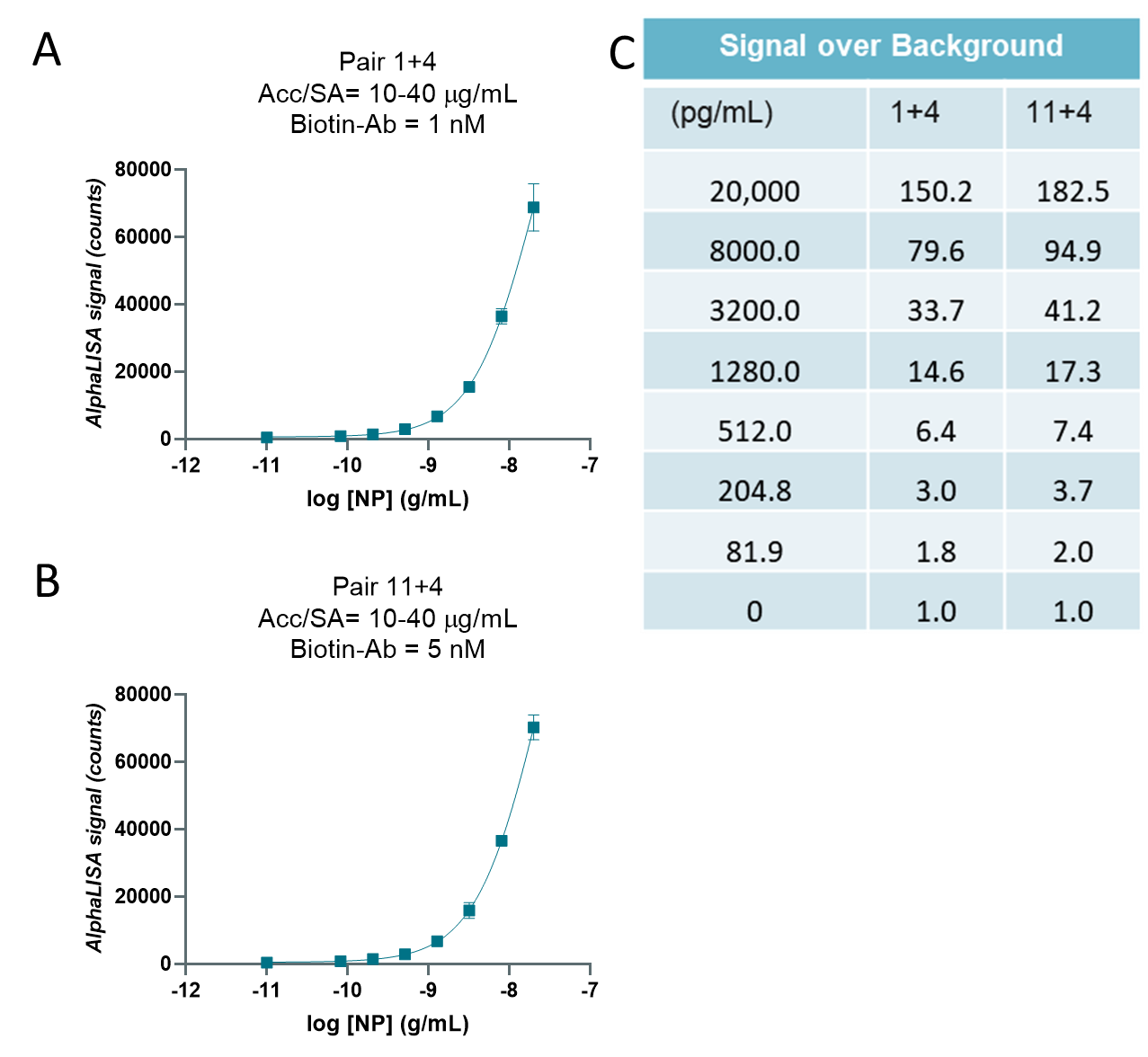

**Supplementary Figure 3.** Standard curves using best conditions for best antibody pairs. **(A)** AlphaLISA signal (counts) for titrated NP (g/mL) from 20,000 pg/mL to 0 pg/mL using ACC/SA = 10-40 µg/mL and 1 nM Biotin-Ab for Pair 1+4. **(A)** AlphaLISA signal (counts) for titrated NP (g/mL) from 20,000 pg/mL to 0 pg/mL using ACC/SA = 10-40 µg/mL and 5 nM Biotin for Pair 11+4. **(C)** Signal/Background table for graphs 1+4 and 11+4. N = duplicate wells. Error bars indicate S.D.

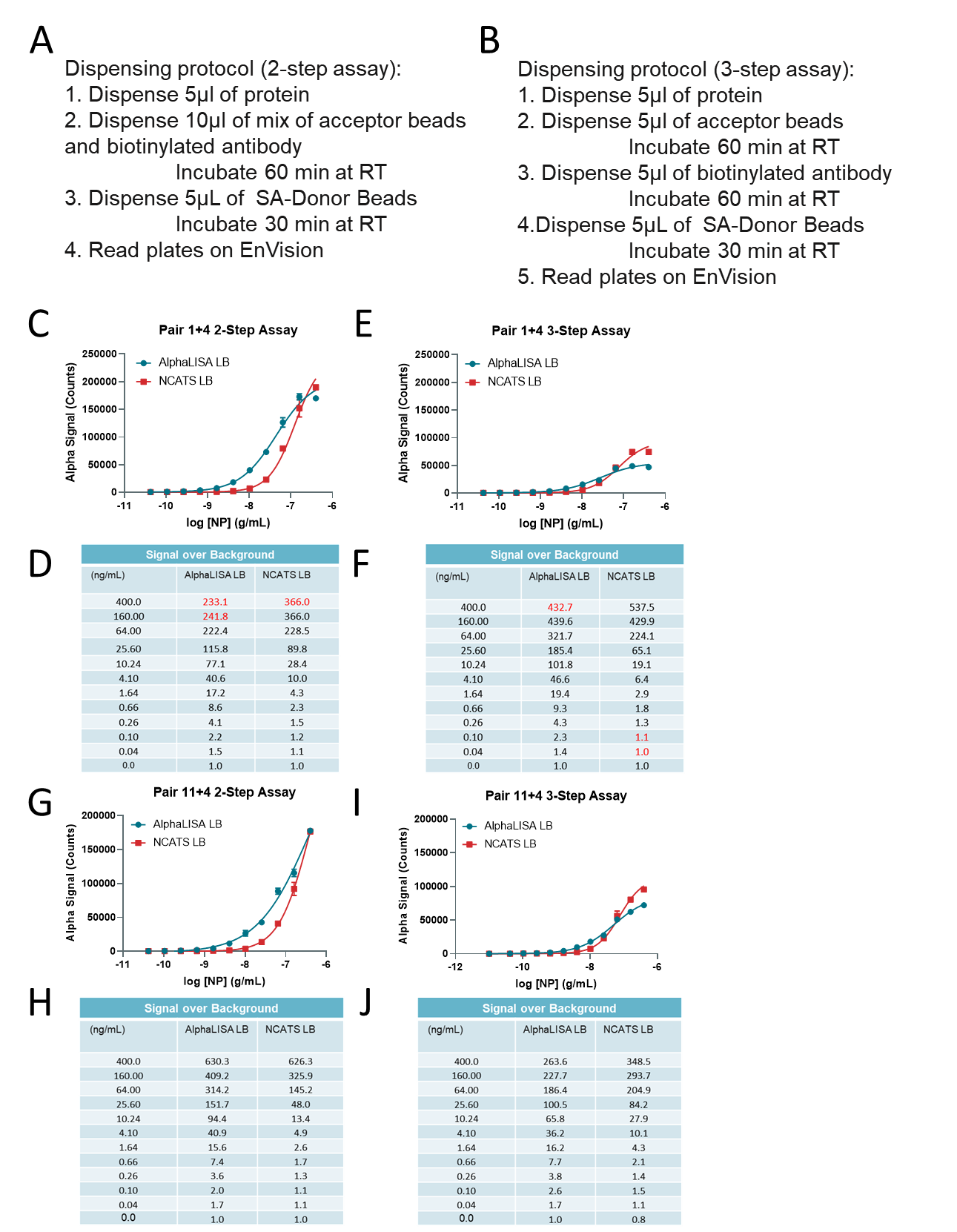

**Supplementary Figure 4** 2-step vs 3-step assay optimization for best antibody pairs and conditions. Dispensing protocols for **(A)** 2-step and **(B)** 3-step assay.	**(C)** Standard curve for NP using Pair 1+4 and two-step assay protocol using AlphaLISA or NCATS LBs. **(D)** S/B table for C. **(E)** Standard curve for NP using Pair 1+4 and three-step assay protocol using AlphaLISA or NCATS LBs. **(F)** S/B table for E. **(G)** Standard curve for NP using Pair 11+4 and two-step assay protocol using AlphaLISA or NCATS LBs. **(H)** S/B table for G. **(I)** Standard curve for NP using Pair 11+4 and three-step assay protocol using AlphaLISA or NCATS LBs. **(J)** S/B table for I. N = triplicate wells. Error bars indicate S.D.

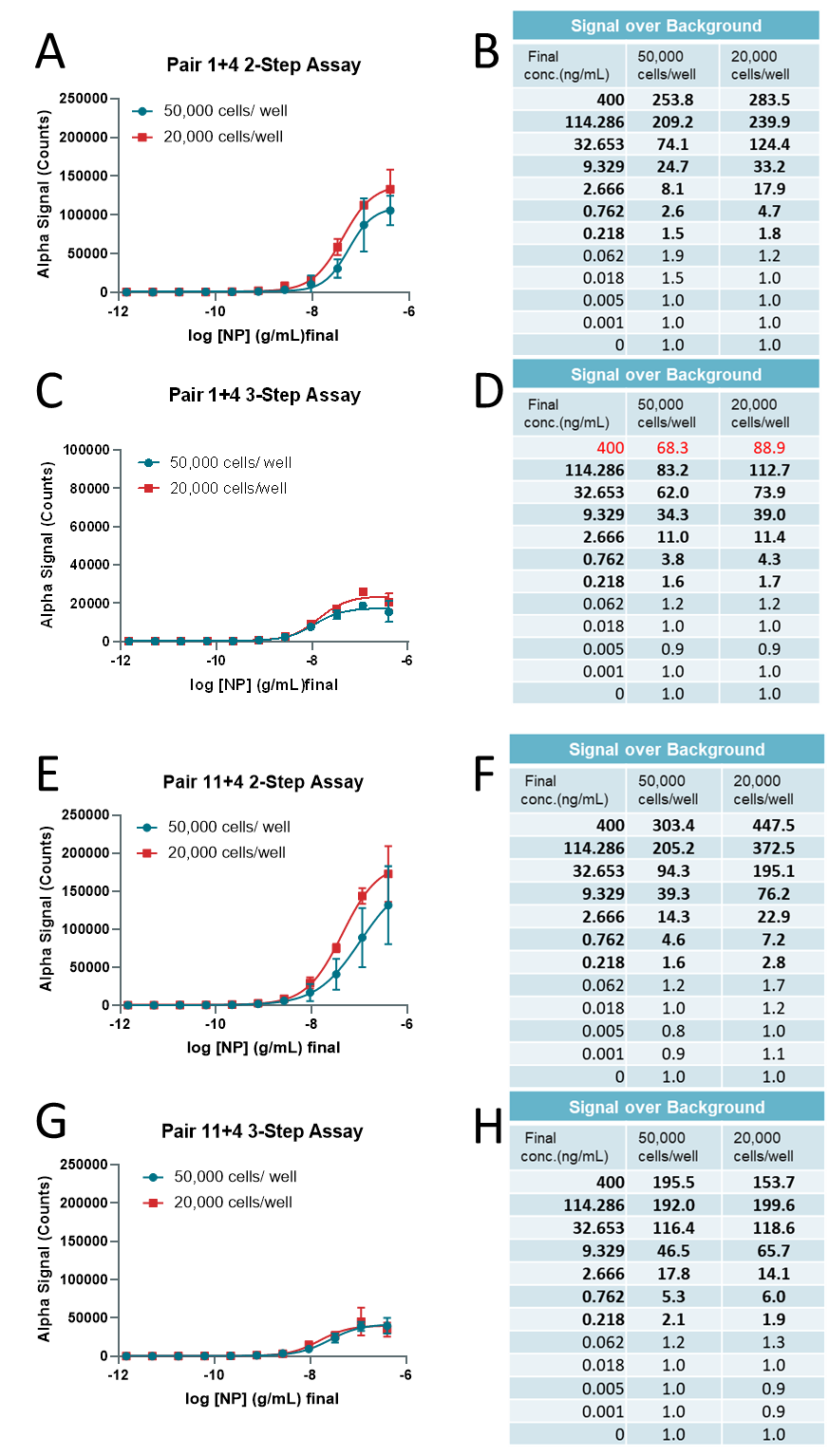

**Supplementary Figure 5** Simulation of viral infection in 384 well plates using recombinant NP spiked into Vero-E6 wells. **(A)** Detection of titrated NP in wells culturing 50,000 or 20,000 Vero-E6 cells using the two-step assay for Pair 1+4. **(B)** S/B table from A. **(C)** Detection of titrated NP in wells culturing 50,000 or 20,000 Vero-E6 cells using the three-step assay for Pair 1+4. **(D)** S/B table from C. **(E)** Detection of titrated NP in wells culturing 50,000 or 20,000 Vero-E6 cells using the two-step assay for Pair 11+4. **(F)** S/B table from E. **(G)** Detection of titrated NP in wells culturing 50,000 or 20,000 Vero-E6 cells using the three-step assay for Pair 11+4. **(H)** S/B table from G. N = triplicate wells. Error bars indicate S.D.

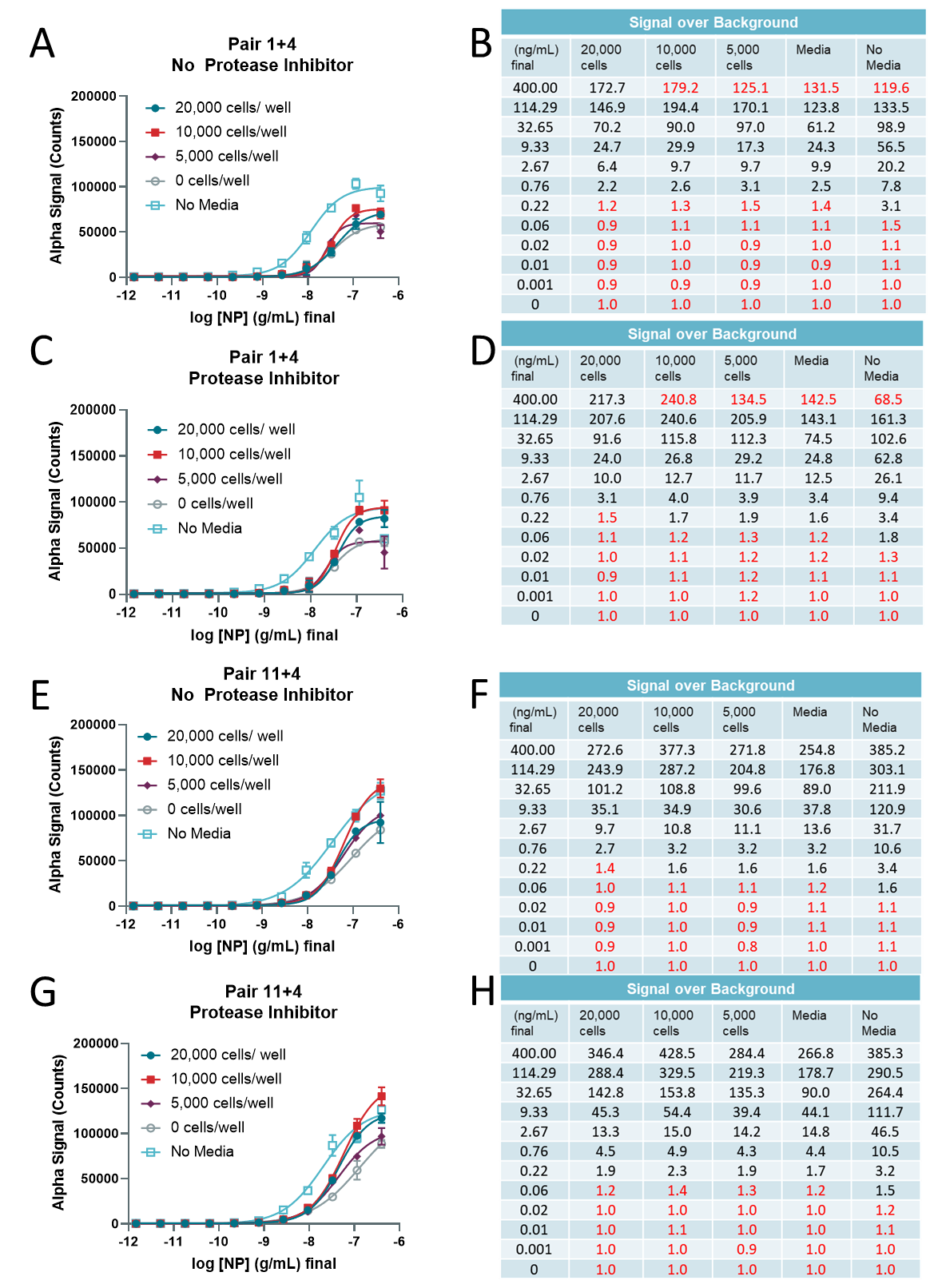

**Supplementary Figure 6** Optimization of LB and cell density for 384-well plate viral infection simulation using recombinant NP. **(A)** Detection of titrated NP in wells culturing 20,000, 10,000, 5000, or 0 cells/well along with a no-media control without protease inhibitor in Vero-E6 cells using Pair 1+4. **(B)** S/B table from A. Red values indicate hook effect. **(C)** Detection of titrated NP in wells culturing 20,000, 10,000, 5000, or 0 cells/well along with a no-media control with protease inhibitor in Vero-E6 cells using Pair 1+4. **(D)** S/B table from C. Red values indicate hook effect. **(E)** Detection of titrated NP in wells culturing 20,000, 10,000, 5000, or 0 cells/well along with a no-media control without protease inhibitor in Vero-E6 cells using Pair 11+4. **(F)** S/B table from E. Red values indicate hook effect. **(G)** Detection of titrated NP in wells culturing 20,000, 10,000, 5000, or 0 cells/well along with a no-media control with protease inhibitor in Vero-E6 cells using Pair 11+4. **(H)** S/B table from G. N = triplicate wells. Error bars indicate S.D.

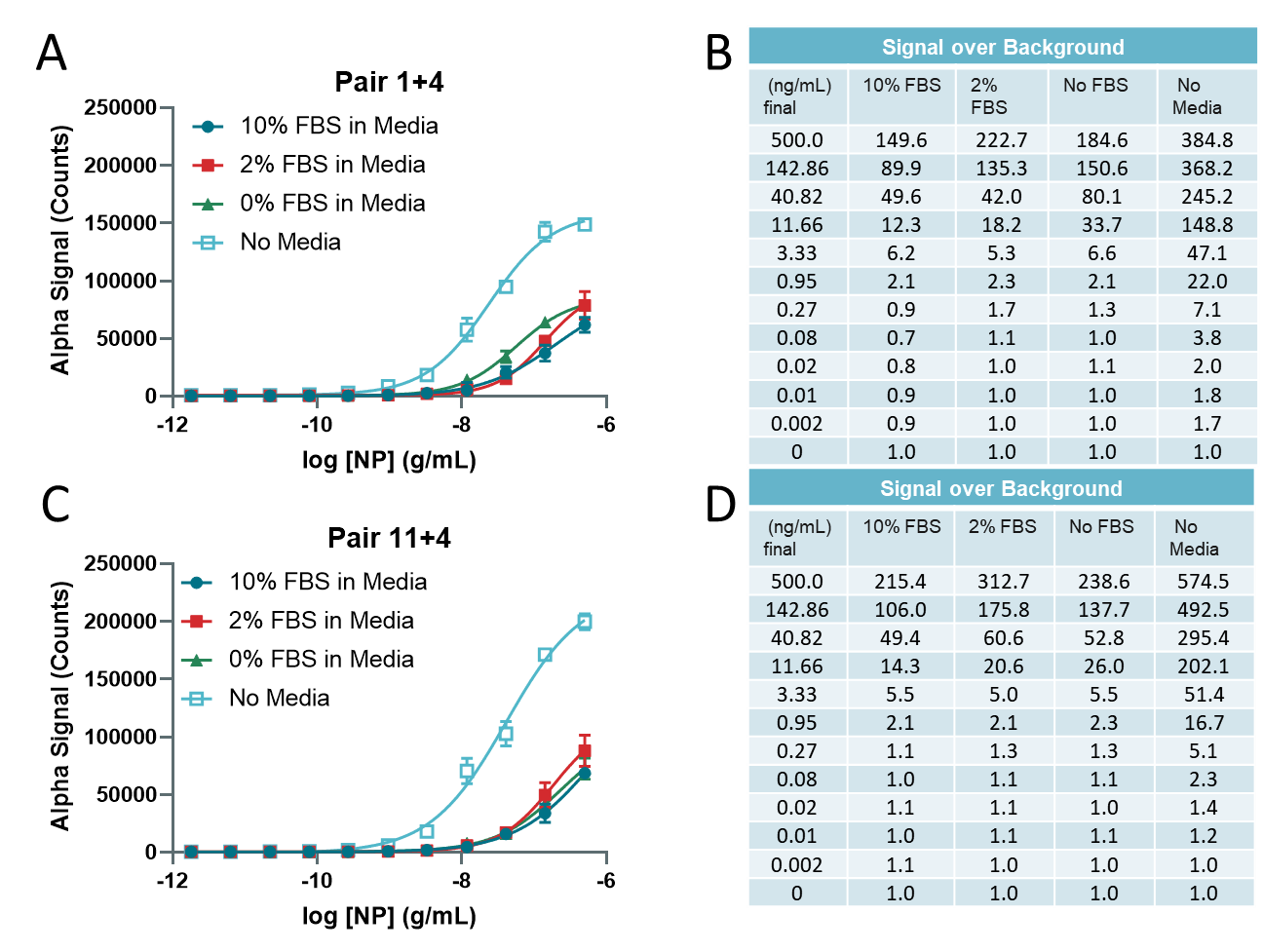

**Supplementary Figure 7** Optimization of AlphaLISA assay using different FBS percentages in cell culture media. **(A)** Detection of titrated NP in wells with 10%, 2% and 0% FBS in media as well as no media control using Pair 1+4. Assay conducted using the AlphaLISA LB. **(B)** S/B table from A. **(C)** Detection of titrated NP in wells with 10%, 2% and 0% FBS in media as well as no media control using Pair 1+4. Assay conducted using the AlphaLISA LB. **(D)** S/B table from A. N = duplicate wells. Error bars indicate S.D.

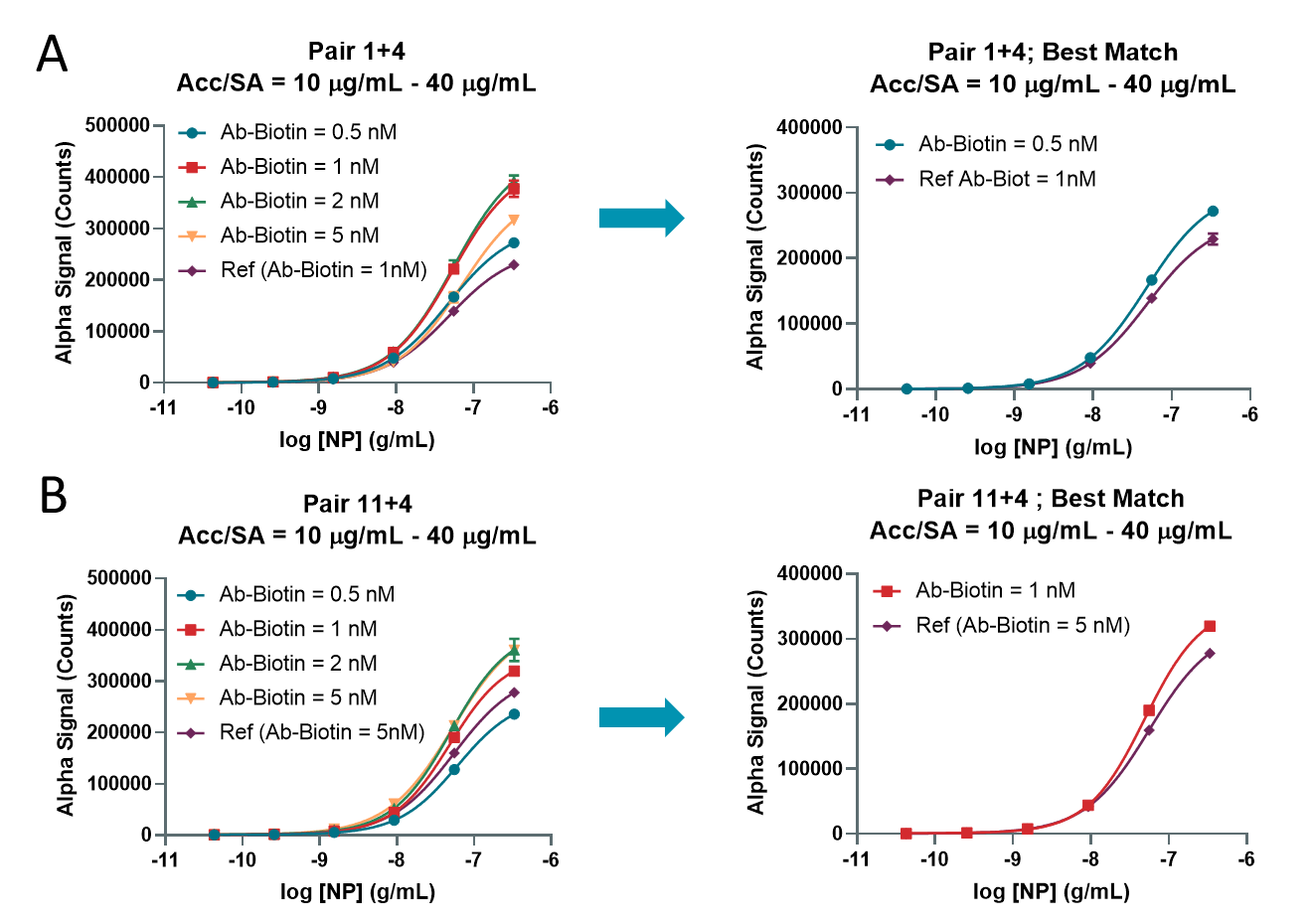

**Supplementary Figure 8** Optimization of best antibody pair and reagent concentrations of second preparation to match first preparation. **(A)** AlphaLISA counts for titrated NP using 10 µg/mL acceptor and 40 µg/mL Streptavidin for Pair 1+4 and different concentrations of Biotin. Right panel comparing the best condition from first batch to matching condition from second batch. **(B)** AlphaLISA counts for titrated NP using 10 µg/mL acceptor and 40 µg/mL Streptavidin for Pair 11+4 and different concentrations of Biotin. Right panel comparing the best condition from first batch to matching condition from second batch. N = duplicate wells. Error bars indicate S.D.

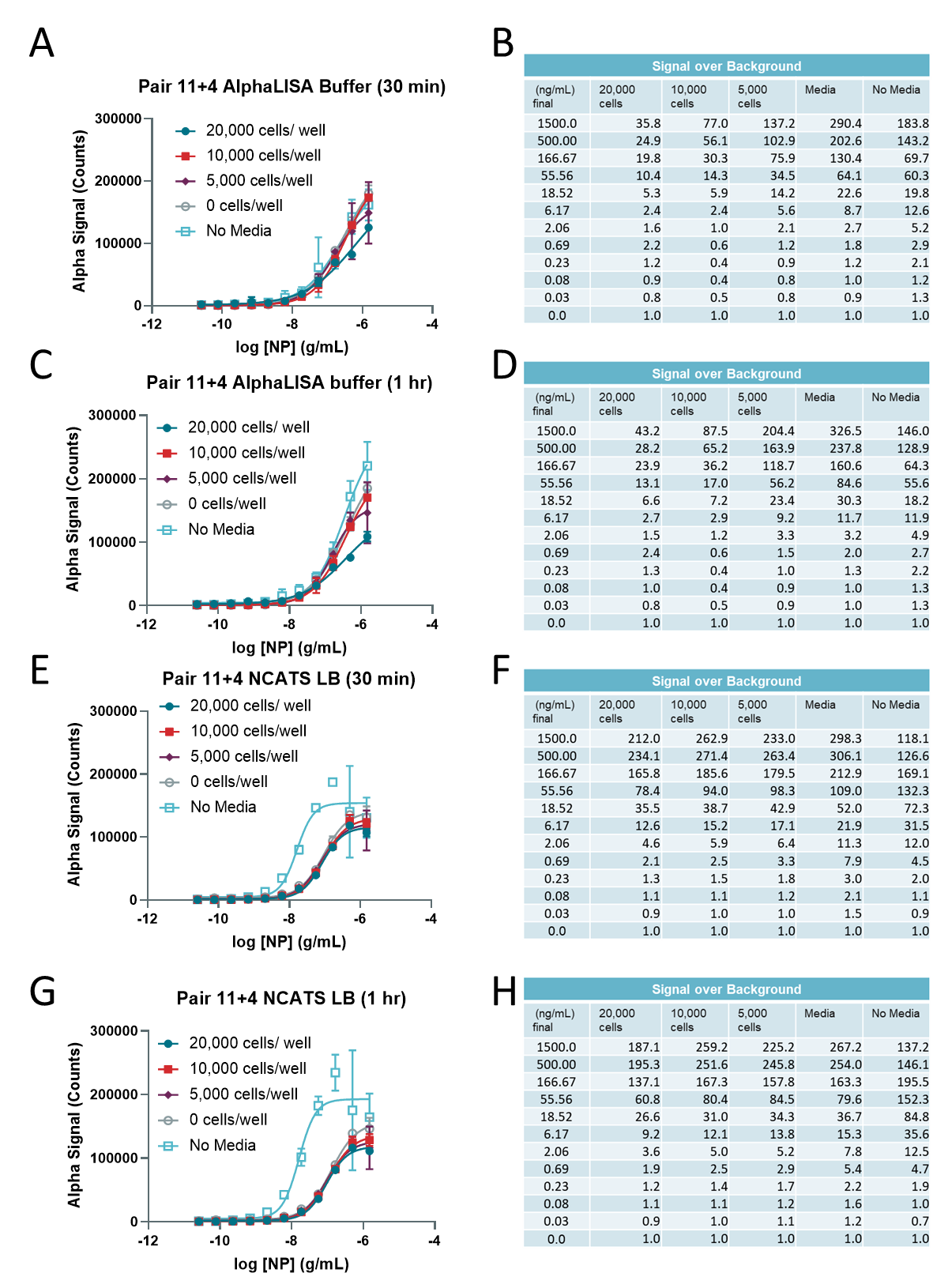

**Supplementary Figure 9** Optimization of LB, cell density, and incubation time for 384-well plate viral infection simulation using recombinant NP and second batch of reagents. **(A)** Detection of titrated NP in wells culturing 20,000, 10,000, 5000, or 0 cells/well along with a no-media control in Vero-E6 cells using Pair 11+4 and AlphaLISA LB after 30 min. **(B)** S/B table from A. Red values indicate hook effect. **(C)** Detection of titrated NP in wells culturing 20,000, 10,000, 5000, or 0 cells/well along with a no-media control in Vero-E6 cells using Pair 11+4 and AlphaLISA buffer after 1 hr. **(D)** S/B table from C. Red values indicate hook effect. **(E)** Detection of titrated NP in wells culturing 20,000, 10,000, 5000, or 0 cells/well along with a no-media control in Vero-E6 cells using Pair 11+4 and NCATS LB after 30 min. **(F)** S/B table from E. Red values indicate hook effect. **(G)** Detection of titrated NP in wells culturing 20,000, 10,000, 5000, or 0 cells/well along with a no media control in Vero-E6 cells using Pair 11+4 after 1 hr. **(H)** S/B table from G. N = triplicate wells. Error bars indicate S.D.

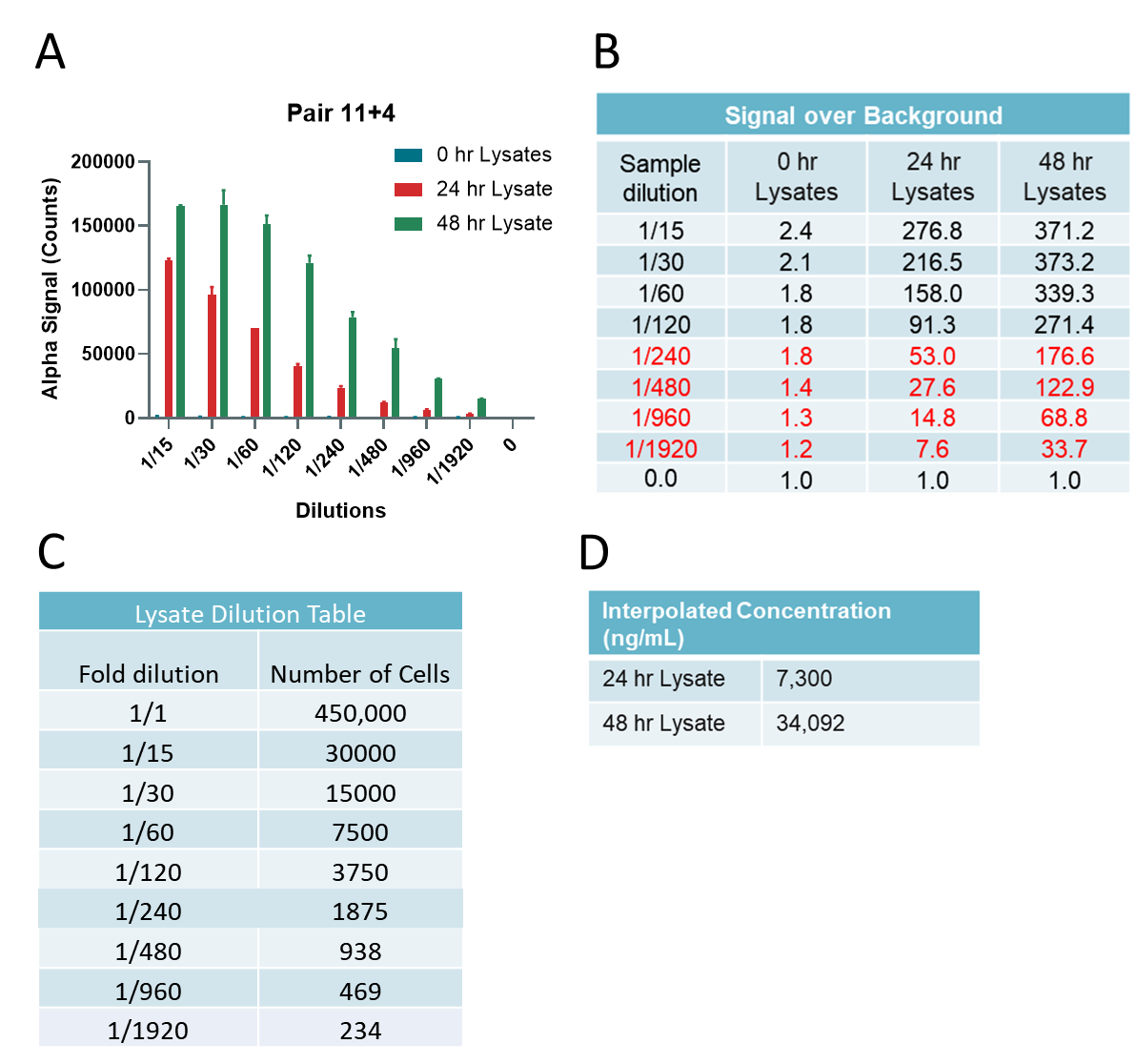

**Supplementary Figure 10** Estimation of viral NP concentration in viral lysates. **(A)** Detection of titrated 0 hr, 24 hr, and 48 hr lysates with NCATS LB using pair 11+4. **(B)** S/B calculation from graph in A. Red values indicate linear range of the curve. **(C)** Interpolated concentrations of NP in 24 hr and 48 hr viral cell lysates. **(D)** Interpolated NP (ng/mL) from samples in A. N = duplicate wells. Error bars indicate S.D.
